# Development of a machine learning model to estimate biotic ligand model-based predicted no□effect concentrations for copper in freshwater

**DOI:** 10.1101/2021.09.16.460690

**Authors:** Jiwoong Chung, Geonwoo Yoo, Jinhee Choi, Jong-Hyeon Lee

## Abstract

The copper biotic ligand model (BLM) has been used for environmental risk assessment by taking into account the bioavailability of copper in freshwater. However, the BLM-based environmental risk of copper has been assessed only in Europe and North America, with monitoring datasets containing all of the BLM input variables. For other areas, it is necessary to apply surrogate tools with reduced data requirements to estimate the BLM-based predicted no-effect concentration (PNEC) from commonly available monitoring datasets. To develop an optimized PNEC estimation model based on an available monitoring dataset, an initial model that considers all BLM variables, a second model that requires variables excluding alkalinity, and a third model using electrical conductivity as a surrogate of the major cations and alkalinity have been proposed. Furthermore, deep neural network (DNN) models have been used to predict the nonlinear relationships between the PNEC (outcome variable) and the required input variables (explanatory variables). The predictive capacity of DNN models in this study was compared with the results of other existing PNEC estimation tools using a look-up table and multiple linear and multivariate polynomial regression methods. Three DNN models, using different input variables, provided better predictions of the copper PNECs compared with the existing tools for four test datasets, i.e., Korean, United States, Swedish, and Belgian freshwaters. The adjusted *r*^*2*^ values in all DNN models were higher than 0.95 in the test datasets, except for the Swedish dataset (adjusted *r*^*2*^ > 0.87). Consequently, the most applicable model among the three DNN models could be selected according to the data availability in the collected monitoring database. Because the most simplified DNN model required only three water quality variables (pH, dissolved organic carbon, and electrical conductivity) as input variables, it is expected that the copper BLM-based risk assessment can be applied to monitoring datasets worldwide.

## 1. Introduction

The copper biotic ligand model (BLM) is used to assess environmental risks and toxicity for copper based on its bioavailability because the toxicity of copper in aquatic systems is highly dependent on site-specific water chemistry. The model assumes that the binding of free copper ions to biotic ligands, together with the competitive effects of major cations, determines copper toxicity [1, 2]. There are a number of essential input variables (pH, dissolved organic carbon (DOC), major cations, and alkalinity) required to derive the predicted no-effect concentration (PNEC) and an effective environmental quality standard based on the copper BLM. However,monitoring databases containing all BLM input variables are available only for a few regions, such as the United States and Europe. Regulatory monitoring databases, which are not intended for use in BLM-based risk assessments, contain only general water quality variables and hazardous substances as monitoring variables.

Although existing PNEC estimation tools can produce uncertain results due to the use of only a few assessment parameters, a BLM-based risk assessment can be conducted in regions where not all of the data required as BLM input variables are available. The Bio-met look-up table, the Environment Agency metal-bioavailability assessment tool (mBAT), which uses a multivariate polynomial function, and PNEC-pro, which uses multiple linear regression (MLR), require pH, DOC, and Ca, as the most influential variables to determine BLM-based PNECs [3-5]. However, the use of Ca as a representative variable of the major cations and alkalinity in existing tools has not significantly broadened the ecoregion for which BLM-based risk assessments can be applied. The Ca content may or may not be included as a common regulatory monitoring variable in different ecoregions. There is a need for new input variables that can act as a surrogate for the major cations and alkalinity within water quality variables while maintaining a good predictive capacity for the BLM-based PNECs. In this study, electrical conductivity was considered a surrogate variable and is one of the recommended variables used to estimate the values of a missing BLM variable [6, 7].

New PNEC estimation models should be developed using a method that minimizes the remaining uncertainty by using the input variables from available monitoring datasets. In this study, a deep neural network (DNN) was used rather than the statistical methods that are applied in existing tools. The DNN was expected to provide an optimized predictive capacity for the nonlinear relationship between the BLM-based PNEC and the BLM input variables. The DNN is an approximator of universal function. It is an artificial neural network consisting of multiple hidden layers between the input and output layers, and therefore complex nonlinear relationships can be modeled by stacking more hidden layers [8].

Another factor determining the predictive capacity of the PNEC estimation model is that the dataset used to develop it must be sufficiently representative of freshwater chemistry. In the dataset used for the development of Bio-met and mBAT, Peters et al. (2011) assumed that most of the Mg, Na, and alkalinity could be determined from Ca concentrations [9]. This means that a dataset consisting of a combination of only three variables (pH, DOC, and Ca) would not cover the full range of BLM input variables. The dataset used for the development of PNEC-pro is from a monitoring database from the Netherlands. Further validation is therefore necessary to apply PNEC-pro to ecoregions with different water chemical properties. As a result, simulation data with full coverage of the domain of BLM input variables is needed for the development of the PNEC estimation model.

The aim of this study was to develop an optimized PNEC estimation model depending on the available monitoring dataset. For this purpose, a realistic training dataset with sufficiently representative freshwater chemistry was built to combine all the BLM input variables, and three different models with a different number of input variables were proposed by the DNN. The most simplified model required only general water quality parameters, such as pH, DOC, and electrical conductivity, and could be used for copper BLM-based risk assessments using various monitoring datasets that are available worldwide.

## 2. Materials and methods

### 2.1. Calculation of the BLM-based PNECs for copper

A general formula for a copper BLM (the *Daphnia magna* BLM) is shown in Eq 1 [10]. According to the European Union Risk Assessment Report (EU-RAR) [11], the acute *D. magna* BLM was used as the chronic fish BLM as follows:

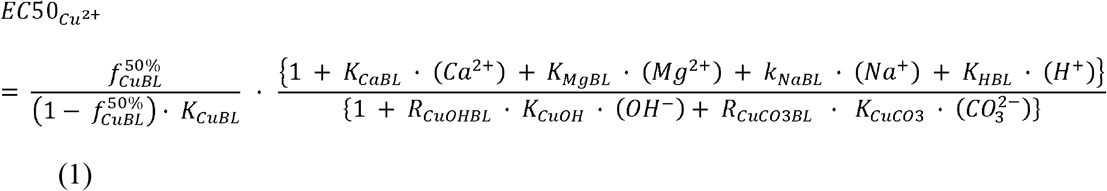

where 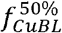 is the fraction of the total number of copper-binding sites occupied by copper at the 50% toxic effect, and K represents biotic ligand constants, such as *K*_*CaBL*_, *K*_*MgBL*_, *K*_*NaBL*_, *K*_*HBL*_, *R*_*CuOHBL*_ (*K*_*CuOH*BL_/ *K*_*Cu*BL_), and *R*_*CuCO3BL*_(*K*_*CuCO*3BL_ / *K*_*Cu*BL_). The formula for the chronic *D. magna* BLM is shown in Eq 2 [12].

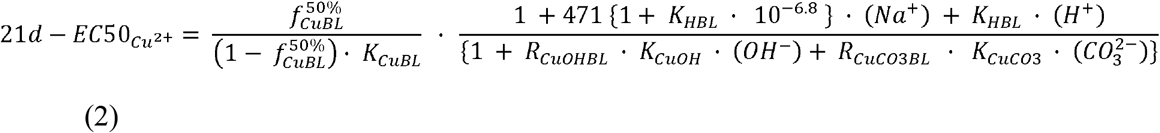

To calculate the BLM-based PNEC in the training and test datasets, site-specific chronic toxicity values were calculated from toxicity data for 27 aquatic organisms provided by the EU-RAR [11]. The biotic ligand and inorganic stability constants for each BLM were applied to three taxonomic groups, algae, invertebrates, and vertebrates, and are shown in S1 Table. The BLM-based PNECs were derived by applying an assessment factor of one to the fifth percentile value (HC5) in the species sensitivity distribution.

### 2.2. Training and test datasets

The training data for DNN model development were built by simulating BLM-based PNECs based on the combination of BLM input variables, including various water chemistry parameters. A monitoring database of Korean freshwater parameters was used to establish the domain range of the training dataset, in which real correlations between BLM input variables were taken into account. The combination of BLM variables was generated from the linear regressions between each variable, and the extent of the domain range was determined by a factor of five of the linear regression results.

Monitoring databases for four ecoregions were used as test datasets. The Korean dataset contained 764 individual samples from the Han River, Guem River, Yeongsan River, and Seomjin River collected from a search of the Environmental Digital Library of the Ministry of Environment from 2014 to 2016 (https://library.me.go.kr). The Swedish dataset contained 4,639 individual samples (999 river samples, 1,914 Malar Lake samples, and 1,726 tributary samples) collected from the Swedish river monitoring program of the Swedish University of Agricultural Sciences from 1997 to 2020 (https://www.slu.se/vatten-miljo). The United States dataset included 279 samples collected in the water monitoring datasets of the Oregon Department of Environmental Quality Water Monitoring Data Portal (https://www.oregon.gov/deq/Data-and-Reports/Pages/default.aspx) and included 84 samples collected from the draft technical support document of the United States Environmental Protection Agency (US EPA, 2016). The Belgian dataset contained 3,187 individual samples reported by Nys et al. (2018) [13].

### 2.3. The DNN models

To estimate the BLM-based PNECs in the available monitoring dataset, an initial model that considered all BLM variables, a second model that required variables excluding alkalinity, and a third model using pH, DOC, and electrical conductivity were developed by a DNN. Optimization of the architecture of the DNN models, which is an artificial neural network composed of several hidden layers between an input layer and an output layer, was performed empirically. The numbers of layers and nodes, which are the main hyperparameters that determine the DNN architecture, were established to minimize the training and validation losses during a fixed period within the search range of hyperparameters, as shown in Table 1. A DNN is generally considered to have at least two hidden layers, and generalization is better with a feedforward neural network with two hidden layers than with one layer according to Thomas et al. (2017) [8]. In this study, the training and validation losses converged to low values when the input layer had three, five, or six nodes, the three hidden layers had 20, 15, or 10 nodes, and the output layer had one node. In addition, these losses decreased stably at a learning rate of 0.005. If the learning rate was 0.1, the losses did not decrease, and if it was less than 0.0001, the losses decreased slowly. The loss values for training were calculated as follows:

**Table 1.** Ranges for deep learning hyperparameter optimization and hyperparameter configuration.

| Hyperparameter | Value | Search range |
| --- | --- | --- |
| Learning rate | 0.005 | 0.1, 0.01, 0.005, 0.001, 0.0005 |
| Optimization method | AdaMax | AdaM, AdaMax, SGD |
| Number of hidden layers | 3 | 1, 2, 3, 5 |
| Number of hidden units | {20, 15, 10} | {20, 15, 10}, {64, 128, 32} |
| Activation functions of hidden layers | {sigmoid, sigmoid, ReLU} | {Sigmoid, Sigmoid, Sigmoid},<br>{Sigmoid, Sigmoid, ReLU},<br>{ReLU, ReLU, Sigmoid},<br>{ReLU, ReLU, ReLU} |
| Batch size | Maximum | Maximum |
| Number of epochs | 20,000 | 500–40,000 |

SGD = stochastic gradient descent; ReLU = rectified linear unit

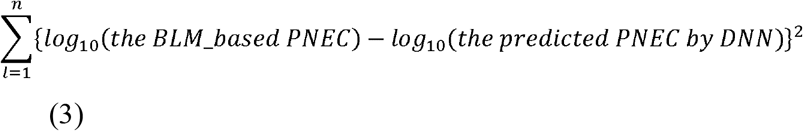

Losses are reduced more by the AdaMax algorithm, which is a variant of the AdaM algorithm based on the infinity norm, than by the AdaM algorithm and the stochastic gradient descent method [14]. The AdaMax algorithm extends the part of the algorithm that adjusts the learning rate based on the L^2^ norm in the AdaM algorithm to the L^p^ norm.

Two different types of activation functions were considered for the DNNs. The sigmoid activation function has traditionally been used as a bounded and monotonically increasing differentiable function. As a remedy for vanishing gradients, the rectified linear unit (ReLU) function [15] has computational advantages over the sigmoid activation function, according to Schmidt-Hieber (2020) [16]. The training and validation losses were reduced more reliably when using the sigmoid function for the first and second hidden layers, and ReLU for the last hidden layer, than when using ReLU for all layers. The epoch, which is the number of iterations of the process of updating the neural network parameters to the loss decreases, was 20,000. For training the dataset, 70% of the randomly shuffled data were used for training and the remaining 30% for validation. The DNN models were implemented using Pytorch version 1.8.1 in Python v3.7 software.

### 2.4. Data Treatment and Statistics

The HC5 for the derivation of PNEC for copper was calculated assuming a log-normal distribution of species sensitivity in the ETX 2.0 software [17]. Normality tests, such as the Anderson–Darling, Kolmogorov–Smirnov, and Cramer von Mises tests, were performed using ETX 2.0 software. A speciation model, such as the Windermere Humic Aqueous Model 7 (WHAM), is required to estimate the site-specific free ion activities for copper and the major cations in training and test datasets [18]. Some element-specific parameters were changed from WHAM-provided values to copper BLM-provided constants (S1 Table). Humic acid and fulvic acid, as input variables of the WHAM, were assumed to be 0.001% and 50% of the DOC concentration, respectively, according to the EU-RAR [11]. The predictive capacity of PNEC estimation tools, including the newly developed DNN models, was compared using the Akaike information criterion (AIC), residual standard error (RSE), and adjusted *r*^*2*^ value. All statistics were calculated using Python v3.5 software.

MLR was performed to determine the appropriate electrical conductivity in the training dataset from the combination of BLM variables, i.e., Ca, Mg, Na, pH, and DOC. The most relevant BLM variables were selected for inclusion in the MLR function for electrical conductivity. The general formula for MLR was as follows:

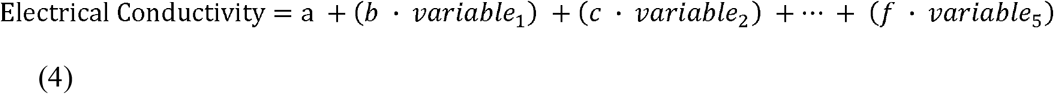

The calculation was completed using a function in R (The R Project for Statistical Computing). Whether the predictive capacity of the MLR model was dependent on the type of BLM variable considered was determined by the AIC [19].

## 3. Results

### 3.1. The development of DNN model for the estimation of the BLM-based PNECs

The DNN models were developed using the training data for the simulated BLM-based PNECs with various combinations of BLM input variables, in which the domain ranges of input variables reflected water chemistry monitoring data from the northern hemisphere. The real correlations among the BLM variables shown in S1 Fig were taken into account to establish the domain range of the training dataset. The extent of these domain ranges was determined by a factor of five of the linear regression results between each variable. The Mg, Na, and K concentrations and alkalinity were generated from the correlations with Ca (Fig 1A). From the combination of these generated variables, only combinations within the domain range were selected to calculate the BLM-based PNEC for copper (Fig 1B). The pH and DOC ranges were 5.5–9.9 and 0.1–50 mg L^-1^, respectively.

**Fig 1.**
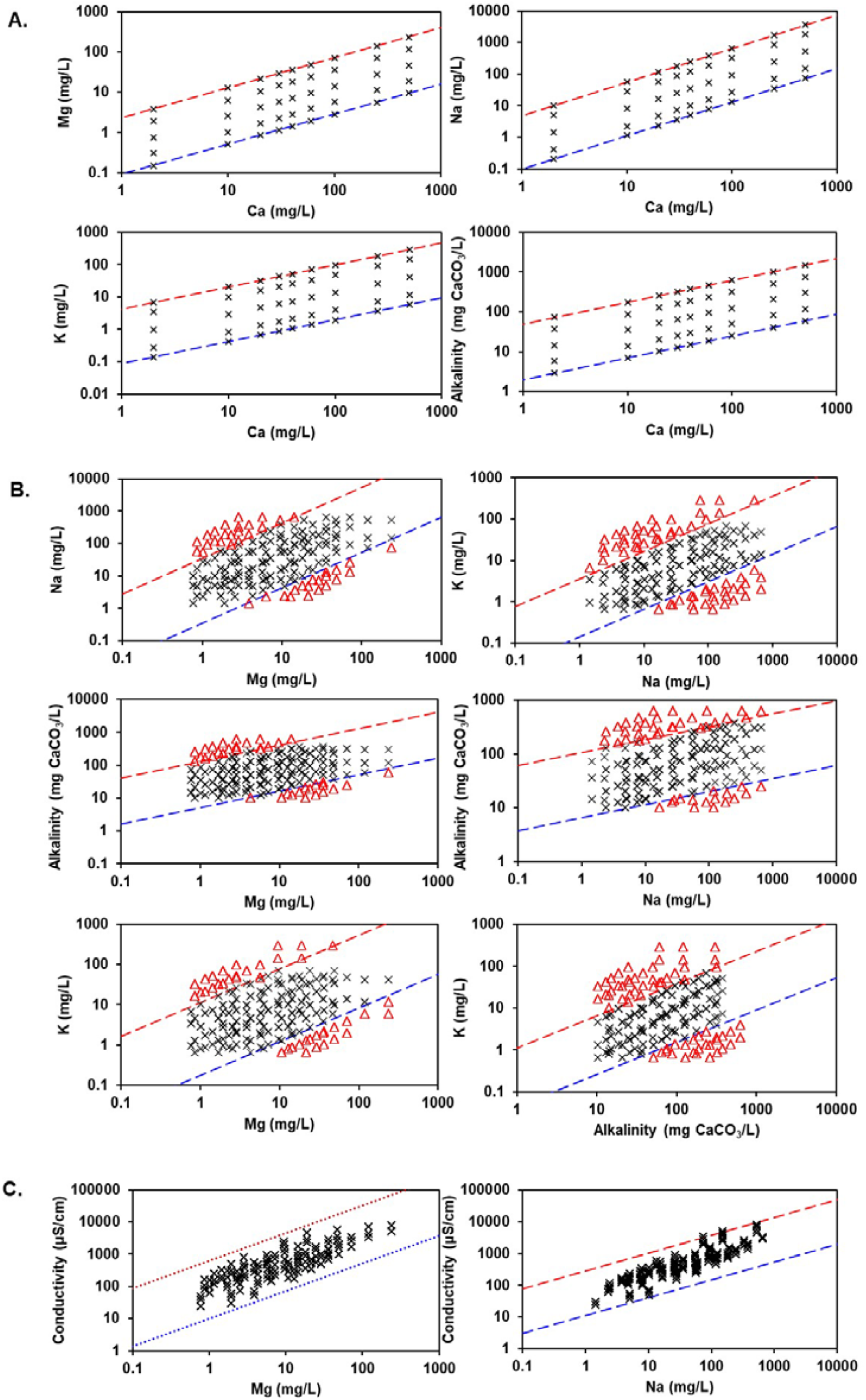
Domain range of input variables in the training dataset used for the development of DNN (deep neural network-based) models as PNEC estimation tools. The dashed lines indicate a factor of five from the linear relationships between variables in Korean freshwaters. The first generated data (cross) are shown in Panel A. The selected data (cross) from the generated data and with the data removed (triangles) outside the domain range are shown in Panel B. The generated electrical conductivity data (cross) added to the selected data are shown in Panel C.

The electrical conductivity estimation model for generating electrical conductivity values from the training dataset was developed by MLR with simplified BLM input variables, using three monitoring datasets (*n* = 5,682) for Korean, Swedish, and the United States freshwaters. Each of the three models required a different number of BLM variables. The first model considered five BLM variables (Ca, Mg, Na, alkalinity, and pH), the second model excluded pH, and the third model excluded pH and alkalinity. The S2 Table shows good agreement between the measured electrical conductivity and the electrical conductivity calculated by the three models (adjusted *r*^*2*^ = 0.959–0.959). As a result, electrical conductivity values in the training dataset were generated using a simplified three-variable (Ca, Mg, and Na) model (Fig 1C).

To develop an optimized PNEC estimation model based on an available monitoring dataset, the DNN(a) model that considered all BLM variables, the DNN(b) model that required all variables excluding alkalinity, and the DNN(c) model that used electrical conductivity as a surrogate of the major cations and alkalinity, were proposed. All of the different DNN models showed a sharp decrease in validation loss after approximately 1,000 epochs without overfitting and flattened out after 10,000 epochs (Fig 2). When the PNECs predicted by the DNN(a), DNN(b), and DNN(c) models within the training dataset were compared with the BLM-based PNECs, the adjusted *r*^*2*^ values were 0.994, 0.990, and 0.965, respectively. As a result, all of the DNN models used in this study were considered sufficiently trained until two constant losses occurred.

**Fig 2.**
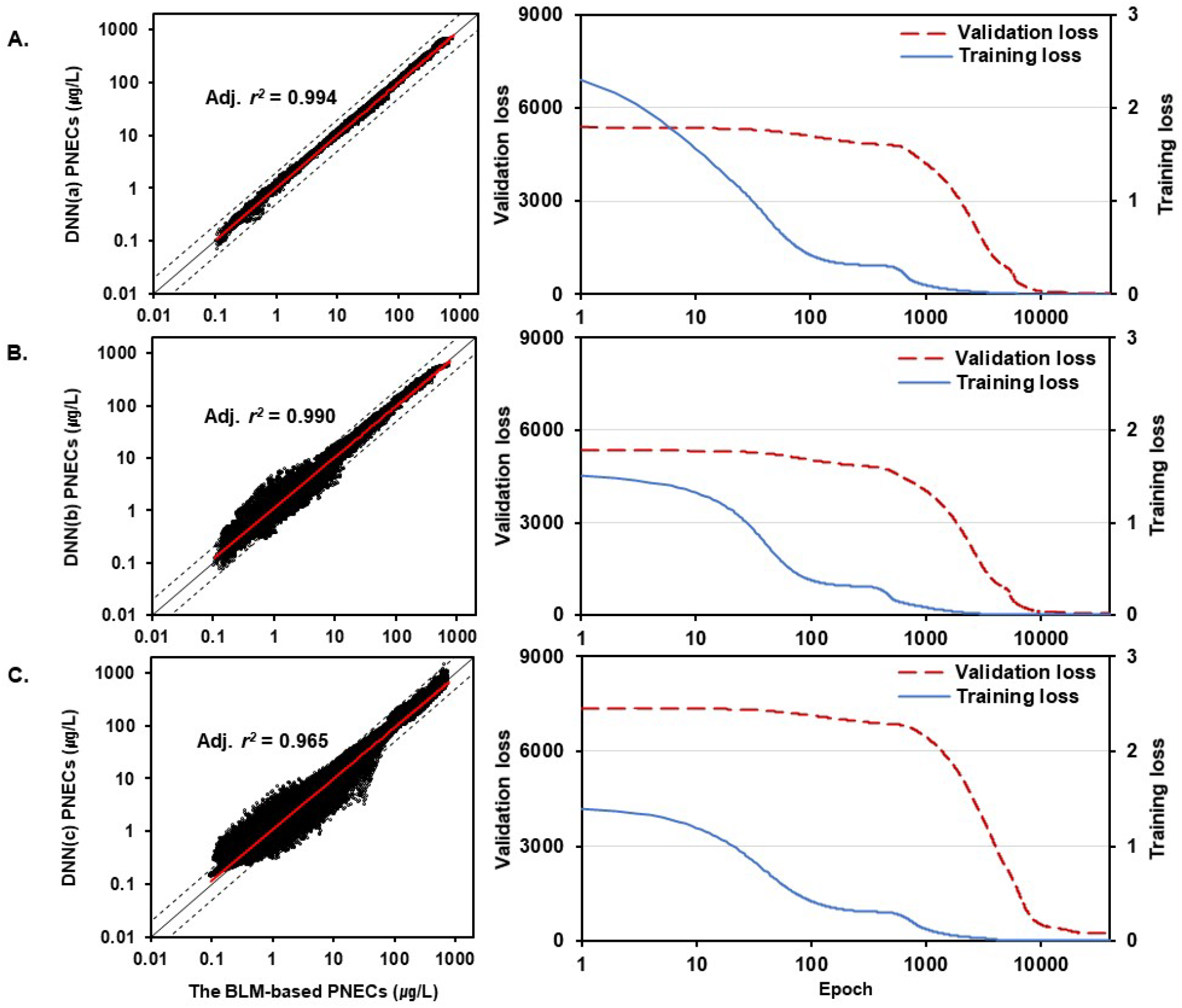
The training and validation results for the DNN(a) model with all BLM variables (A), DNN(b) with all BLM variables except alkalinity (B), and DNN(c) with the three variables of pH, DOC, and electrical conductivity (C). The average loss per epoch for the training and validation steps is shown in the right panels. The validation for the three different types of DNN models within the training dataset is shown in the left panels. The blue solid line indicates loss per epoch for training steps, and the red dashed line indicates loss per epoch for validation steps. The black solid line indicates a perfect match between the simulated and predicted BLM-based PNECs. The black dotted line indicates an error of a factor of two between simulated and predicted BLM-based PNECs. Adj. *r*^*2*^ = adjusted *r*^*2*^ value.

### 3.2. Comparison of PNEC estimation tools with newly developed DNN models

The four test datasets, Korean, United States, Belgian, and Swedish freshwaters, were used to evaluate the predictive capacity of the DNN models and the existing PNEC estimation tools. The differences in water chemistry properties among these four test datasets are shown in S2 Fig as a histogram of the frequency versus concentration of each variable. Korean freshwater had the lowest Ca and DOC concentrations (95^th^ percentile: 16 mg Ca L^-1^ and 8.5 mg DOC L^-1^) and the highest pH (95^th^ percentile: 8.9). Swedish freshwater had the lowest sodium concentration (95^th^ percentile: 26 mg Na L^-1^), and Belgian freshwater had the lowest alkalinity (95^th^ percentile: 13 mg CaCO_3_ L^-1^). United States freshwater had the highest alkalinity (95^th^ percentile: 169 mg CaCO_3_ L^-1^). The application coverage of the DNN model for various water chemistry conditions was dependent on the range of variables in the simulated training dataset. This dataset was considered to be more broadly representative of the water chemistry range compared with the test datasets, and these results affected the predictive capacity of the DNN models (Fig 3).

**Fig 3.**
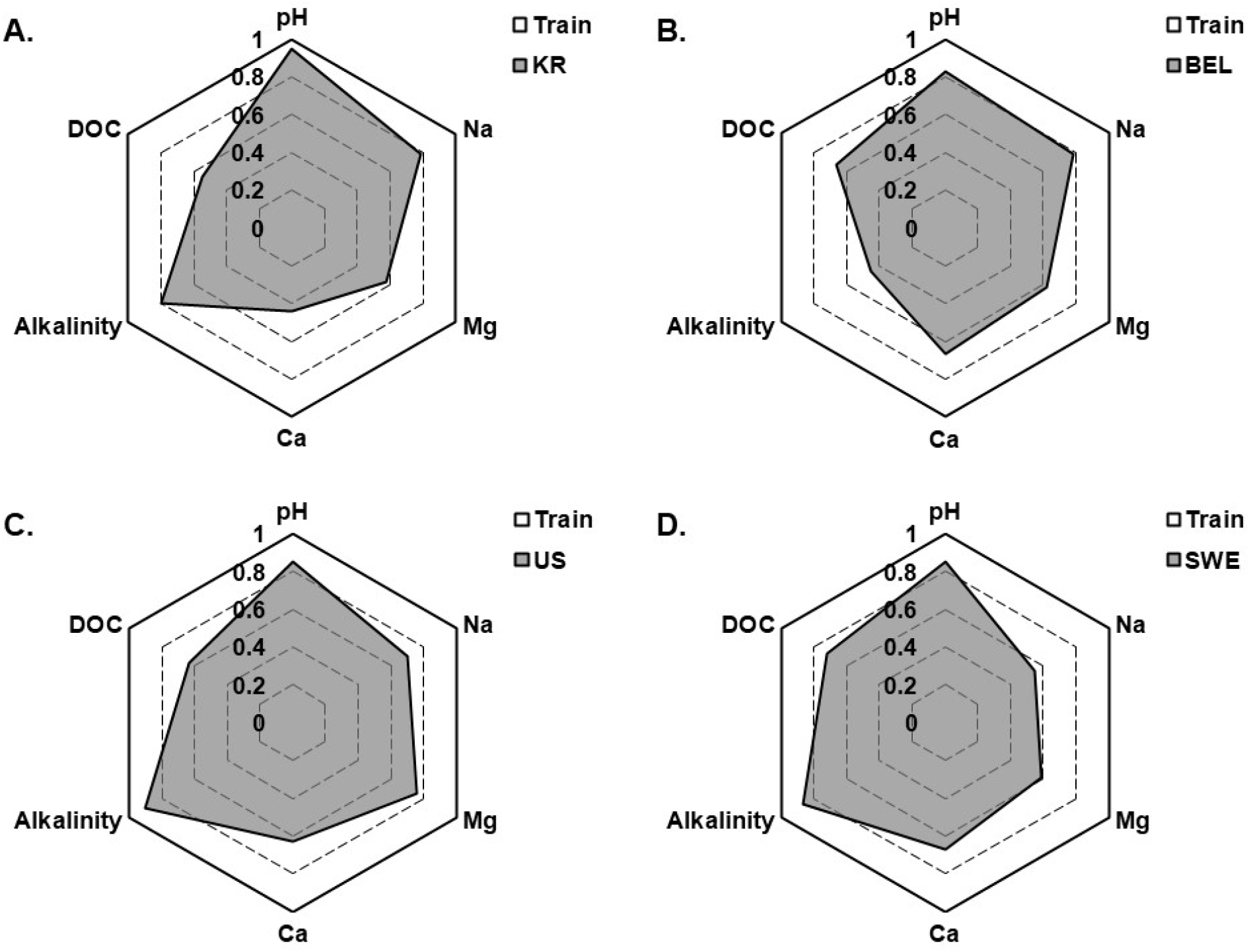
Radar chart showing the ratios of pH and log10 values (different BLM input variables) of four different test datasets to those of the training dataset. The BLM input variables in the training dataset are marked by light shading. The BLM variable ratios in the test datasets are marked as the 95^th^ percentile within the test datasets by dark shading. Train = training dataset; KR = Korean freshwater; BEL = Belgian freshwater; US = United States freshwater; SWE = Swedish freshwater.

Evaluation of the predictive capacity of the three DNN models in this study and comparison of the results with those obtained by existing tools were performed for four ecoregions (test datasets), and the results are shown in Table 2. For Korean freshwater, comparison of the predictive capacity among the PNEC estimation models is shown in Fig 4. The DNN(a) model provided good predictions (adjusted *r*^2^ = 0.987, *p* < 0.01). The DNN(b) and DNN(c) models provided predictions similar to those of DNN(a) (adjusted *r*^2^ = 0.968 and 0.978, respectively, *p* < 0.01). Among the existing models, PNEC-pro provided less reliable predictions (adjusted *r*^2^ = 0.537, *p* < 0.05), whereas Bio-met and mBAT provided good predictions (adjusted *r*^2^ = 0.904 and 0.937, *p* < 0.01).

**Table 2.**
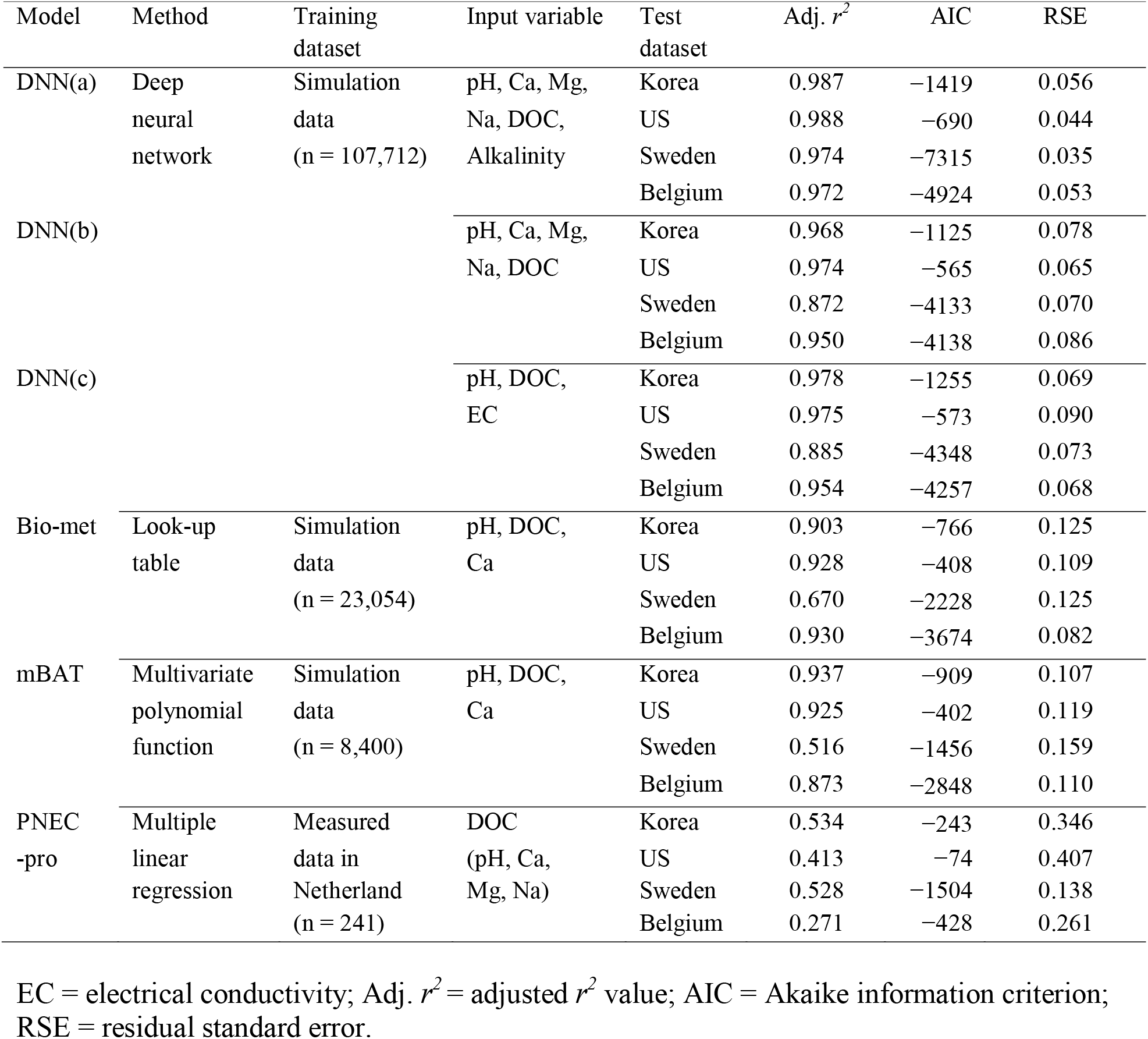
Comparison of newly developed deep neural network models with the existing predicted no-effect concentration estimation tools

**Fig 4.**
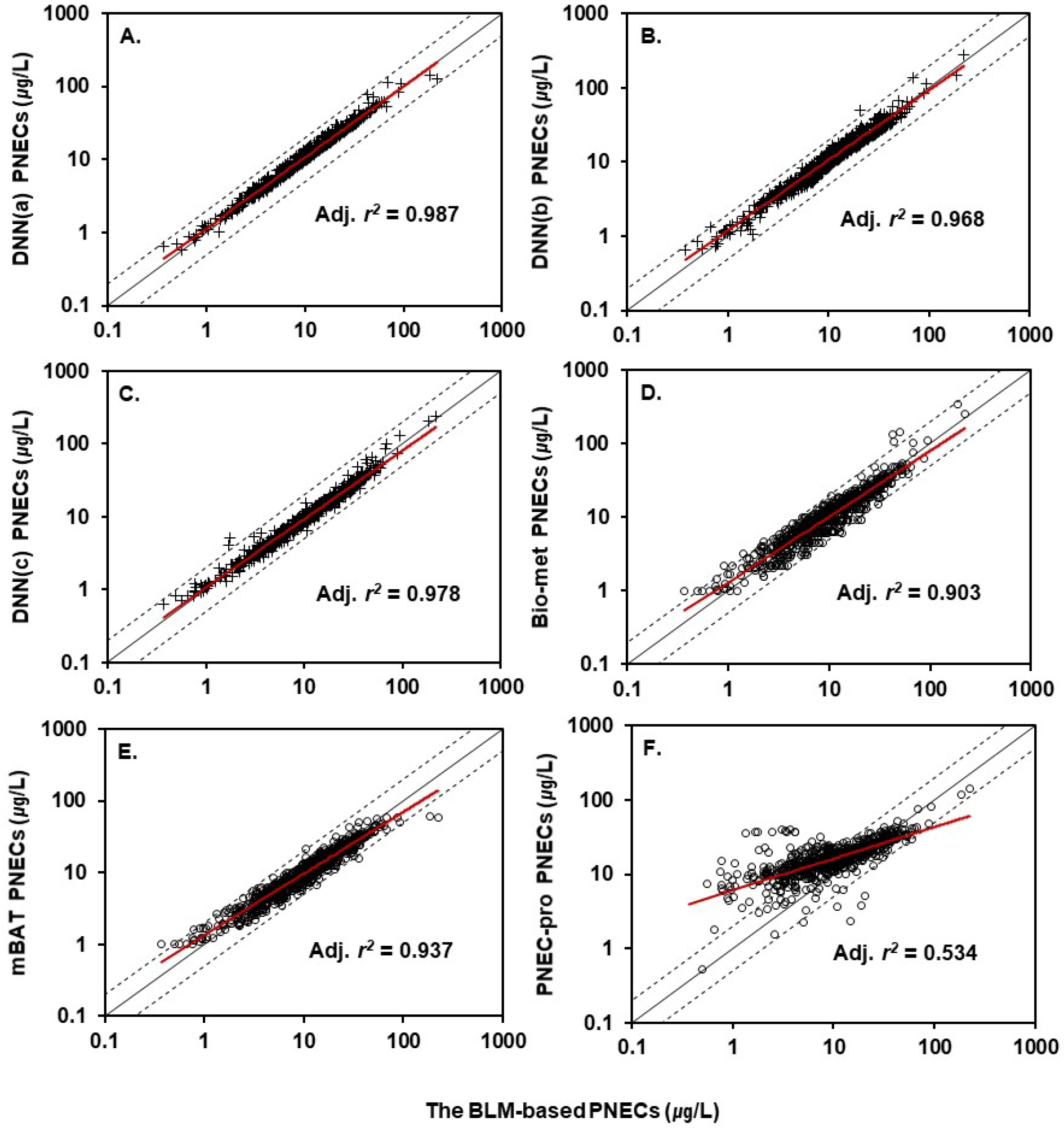
Comparison of the test results of the surrogate models for copper BLM-based PNECs in Korean freshwater. The BLM-based PNECs were derived from 764 individual samples collected in 2014, 2015, and 2016. Panels A, B, and C show PNECs (plus) estimated by the deep neural network-based models DNN(a), DNN(b), and DNN(c), respectively. Panels D, E, and F show PNECs (open circle) estimated by Bio-met, mBAT, and PNEC-pro, respectively. Adj. *r*^*2*^ = adjusted *r*^*2*^ value.

For Swedish freshwater, a comparison of the predictive capacity between the PNEC estimation models is shown in Fig 5. The DNN(a) model also provided good predictions (adjusted *r*^2^ = 0.974, *p* < 0.01). The coefficients of determination of the DNN(b) and DNN(c) models were similar (adjusted *r*^2^ = 0.872 and 0.885, respectively, *p* < 0.01), and were lower than those of DNN(a). For the existing models, the coefficients of determination were lower than 0.7 (adjusted *r*^2^ = 0.670 for Bio-met, 0.529 for PNEC-pro, and 0.516 for mBAT, *p* < 0.05).

**Fig 5.**
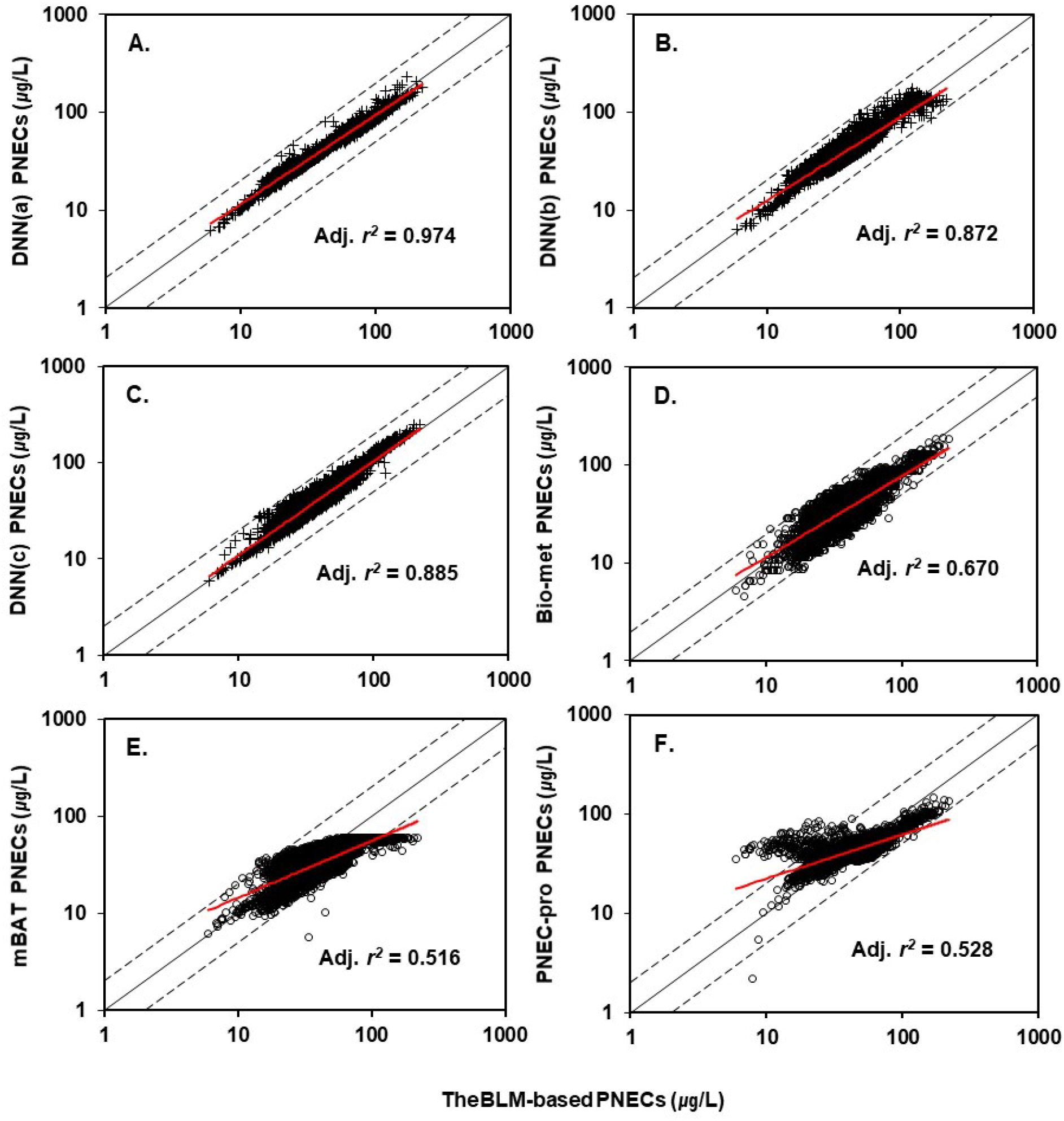
Comparison of the test results of the surrogate models for copper BLM-based PNECs in Swedish freshwater. The BLM-based PNECs were derived from 4,639 individual samples (999 river samples, 1,914 Malar Lake samples, and 1,726 tributary samples) collected in the Swedish river monitoring program of the Swedish University of Agricultural Sciences from 1997 to 2020. Panels A, B, and C show PNECs (plus) estimated by deep neural network-based DNN(a), DNN(b), and DNN(c), respectively. Panels D, E, and F show PNECs (open circle) estimated by Bio-met, mBAT, and PNEC-pro, respectively. Adj. *r*^*2*^ = adjusted *r*^*2*^ value.

For United States freshwater, a comparison of the predictive capacity among the PNEC estimation models is shown in Fig 6. The three DNN models provided good predictions (adjusted *r*^2^ = 0.989 for DNN(a), 0.974 for DNN(b), and 0.975 for DNN(c), *p* < 0.01). Among the existing tools, Bio-met and mBAT provided good predictions (adjusted *r*^2^ = 0.929 and 0.926, respectively, *p* < 0.01), whereas PNEC-pro provided less reliable predictions (adjusted *r*^2^ = 0.421, *p* < 0.05). For Belgian freshwater, a comparison of the predictive capacity among the PNEC estimation models is shown in Fig 7. The coefficients of determination of the three DNN models and Bio-met were > 0.9 (adjusted *r*^2^ = 0.972 for DNN(a), 0.95 for DNN(b), 0.954 for DNN(c), and 0.93 for Bio-met, *p* < 0.01). The mBAT also provided good predictions (adjusted *r*^2^ = 0.873, *p* < 0.01), whereas PNEC-pro provided less reliable predictions (adjusted *r*^2^ = 0.273, *p* < 0.05).

**Fig 6.**
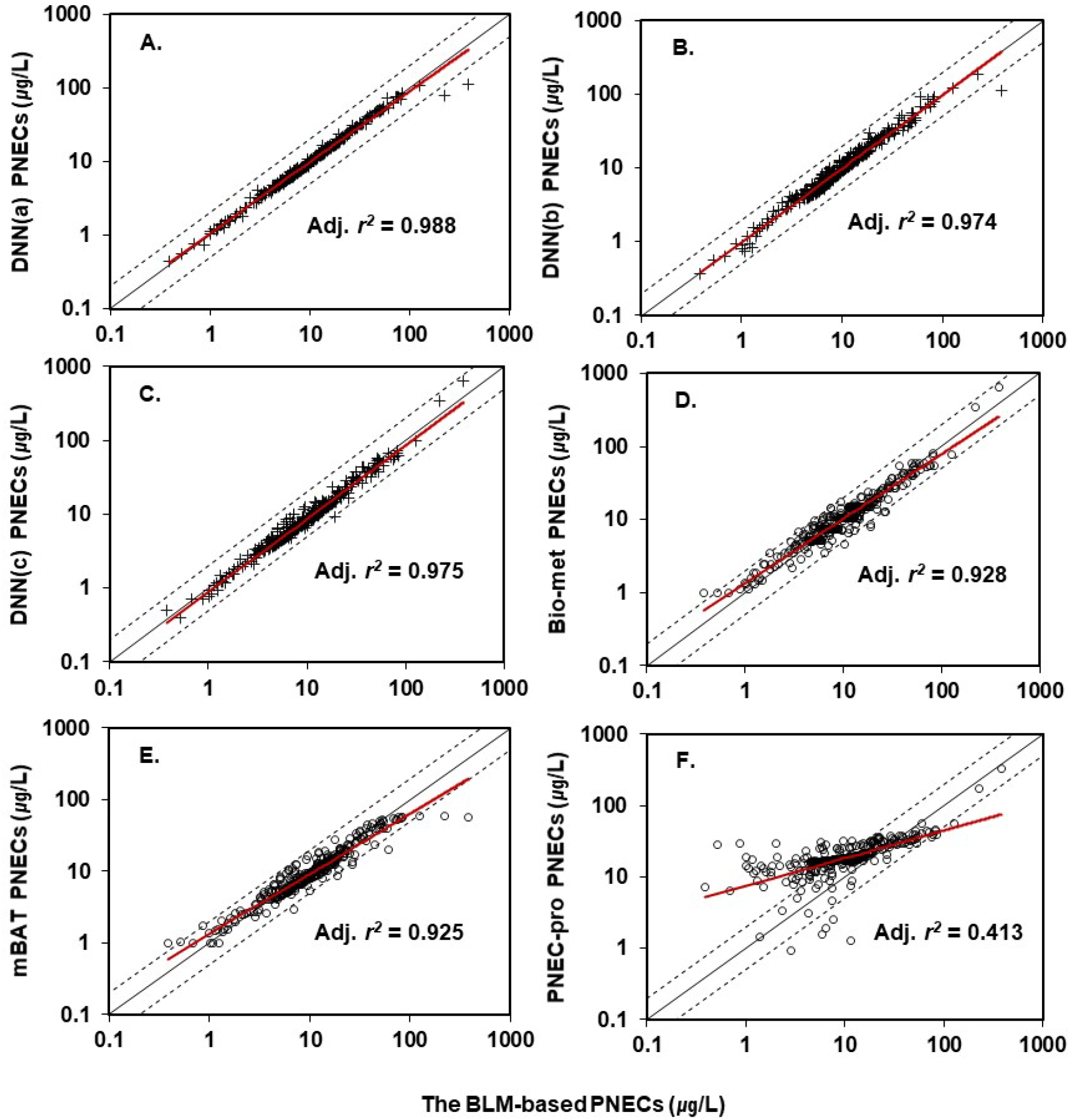
Comparison of the test results of the surrogate models for copper BLM-based PNECs in United States freshwater. The BLM-based PNECs were derived from 363 samples collected by the Oregon Department of Environmental Quality Water Monitoring Data Portal and the National Waters Information System. Panels A, B, and C show PNECs (plus) estimated by the deep neural network-based models DNN(a), DNN(b), and DNN(c), respectively. Panels D, E, and F show PNECs (open circle) estimated by Bio-met, mBAT, and PNEC-pro, respectively. Adj. *r*^*2*^ = adjusted *r*^*2*^ value.

**Fig 7.**
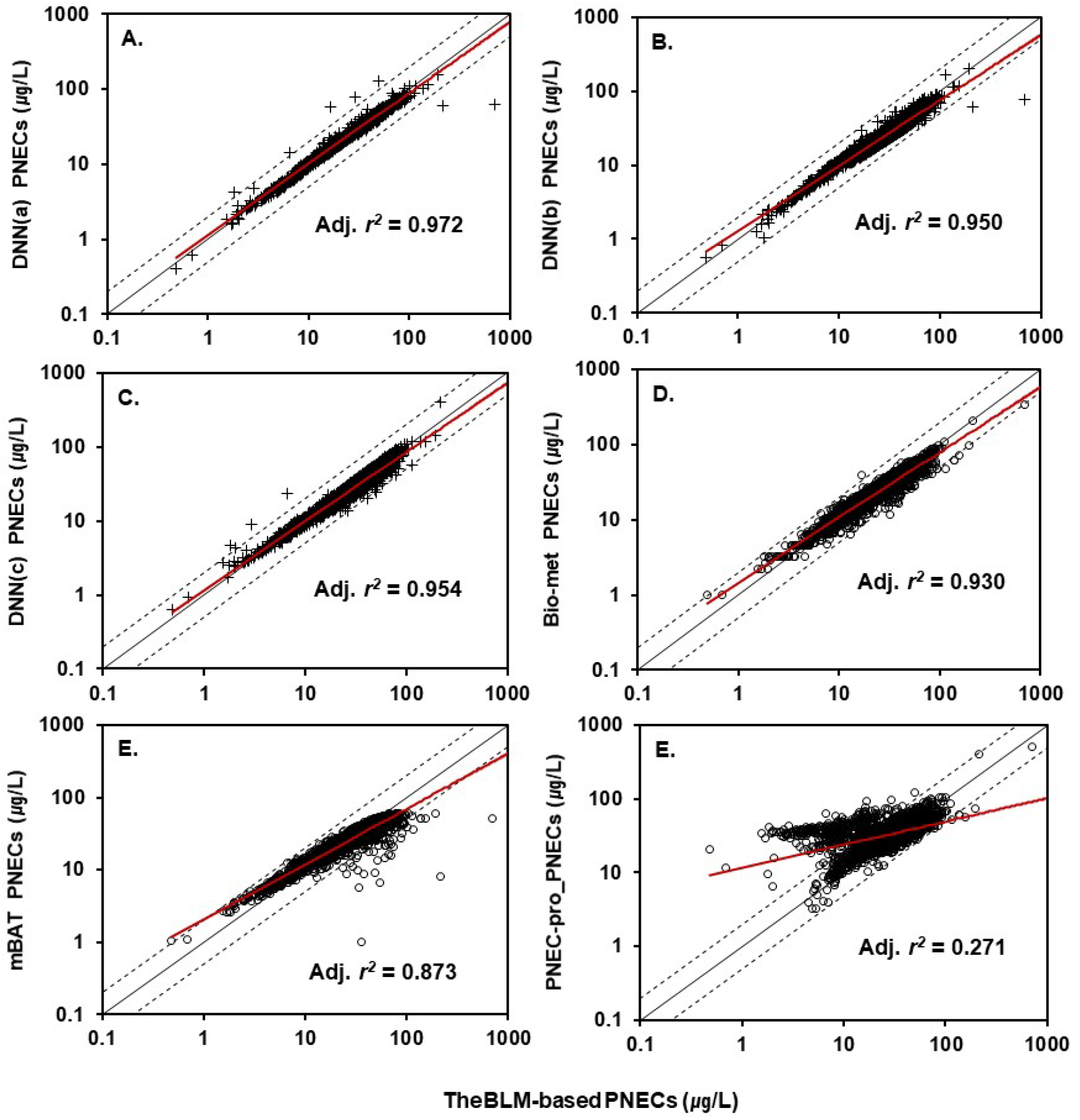
Comparison of the test results of the surrogate models for copper BLM-based PNECs in Belgian freshwater. The BLM-based PNECs were derived from 3,187 individual samples collected by Nys et al. (2018). Panels A, B, and C show PNECs (plus) estimated by the deep neural network-based models DNN(a), DNN(b), and DNN(c), respectively. Panels D, E, and F show PNECs (open circle) estimated by Bio-met, mBAT, and PNEC-pro, respectively. Adj. *r*^*2*^ = adjusted *r*^*2*^ value.

Consequently, all PNEC estimation models based on the DNN method provided good predictions in the four ecoregions (Table 2). The DNN(a) model using all BLM input variables had the lowest AIC and RSE values and the highest adjusted *r*^2^. The DNN(c) model using the variables of electrical conductivity, pH, and DOC had the second lowest AIC and RSE values and the second highest adjusted *r*^2^. The DNN(b) model using five BLM variables (excluding alkalinity) also provided good predictions, which were very similar to those of DNN(c).

Among the existing PNEC estimation tools, the lowest AIC and highest adjusted *r*^2^ values were obtained for Bio-met, based on the look-up table method, while the second lowest AIC and second highest adjusted *r*^2^ were obtained for mBAT, based on a multivariate polynomial function with interaction terms. Compared with the other models, PNEC-pro, based on MLR, had a less reliable predictive capacity for the test datasets.

## 4. Discussion

### 4.1. The development of DNN model for the estimation of the BLM-based PNECs

To develop an optimized PNEC estimation model based on available monitoring datasets, the DNN(a) model that considered all BLM variables, the DNN(b) model that required all variables excluding alkalinity, and the DNN(c) model that used electrical conductivity as a surrogate of the major cations and alkalinity were proposed. These three types of BLM-based PNEC estimation models, using training dataset with various water chemistries, were developed by a DNN to optimize the prediction of nonlinear relationships between input variables (explanatory variables) and BLM-based PNECs (dependent variables). The learning result of the DNN(a) model was predicted to be within a factor of two of that of the BLM-based PNEC for 100% of the data in the training dataset (*n* = 107,712) (Fig 2). This was an expected result because the DNN used for model development was a universal approximation function and was the result of the excellent learning of nonlinear relationships based on large amounts of simulated data. Because simulation data with full coverage of the domain of input variables were used as the training dataset, there was no need to use additional validation and test datasets. The learning results of the DNN(b) and DNN(c) models were predicted to be within a factor of two of the BLM-based PNECs for 98.5% and 88.3% of the data, respectively.

Among the existing PNEC estimation tools, mBAT was developed using a multivariate polynomial function to predict the nonlinear relationships between input variables (pH, DOC, and Ca) and the BLM-based PNECs for copper [4]. Although two functions were proposed for Ca (> and < 6 mg L^-1^) to counteract low Ca concentrations, the validation results of the prediction accuracy for PNECs within the dataset used for development have not been described. PNEC-pro was developed by a simple MLR using monitoring data (*n* = 241) from the Netherlands and provides validation results for the prediction accuracy (adjusted *r*^*2*^ = 0.882) [5]. After determining the MLR function from the learning data of this study, the validation results are shown in S3 Fig. The adjusted *r*^*2*^ value was 0.838, which was lower than that of the DNN models (adjusted *r*^*2*^ = 0.965 for DNN(c) using three variables, Fig 2C). As a result, the DNN models including the most simplified model can be considered the most appropriate method to optimize the prediction of the nonlinear relationship between the required input variables and the BLM-based PNECs in a large training dataset reflecting water chemistry monitoring data from the northern hemisphere.

### 4.2. Comparison of existing PNEC estimation tools with newly developed DNN models

A copper BLM-based PNEC has been proposed in Europe and the United States for environmental risk assessment, taking into account the site-specific bioavailability of copper [11, 19]. To derive the BLM-based PNEC, monitoring datasets including all BLM input variables (pH, DOC, major cations, and alkalinity) are essential for estimating water chemistry speciation, such as the activity of free copper ions, copper speciation, and major cations. However, these datasets are available only in a few regions, such as the United States and Europe. Because some BLM variables may be missing from available datasets, several methods have been proposed to estimate the values of the missing variables [6, 9].

To simulate the derivation of BLM-based PNECs that require all of these input variables, simplified and user-friendly PNEC estimation tools using a reduced number of variables (e.g., Bio-met, mBAT, and PNEC-pro) have been proposed [3-5]. Among these tools, the minimum data requirements for Bio-met and mBAT are pH, DOC, and Ca. pH affects copper toxicity in aquatic organisms and is routinely measured in field samples using a variety of water quality measurement instruments. DOC in freshwater can bind copper and reduce the interaction between free copper ions and aquatic organisms. Non-linear relationships among pH, copper toxicity, and the binding properties of DOC have been reported in EU-RARs [11]. Although the Ca concentration or hardness is a less influential variable than pH and DOC, it is a more statistically effective variable for PNEC than other cations and alkalinity [5]. In addition, it has been reported that an increase in the Ca concentration does not result in an increase in PNEC [9]. However, it may or may not be included as a general water quality variable in regulatory monitoring databases. Therefore, Bio-met and mBAT, which only require the concentration of Ca among the major cations, do not significantly broaden the ecoregion where a BLM-based risk assessment can be applied. Because Ca, Mg, and Na are monitoring variables that can be measured by the same analyzer in one sample, it may be more efficient to improve the predictive capacity by using the concentrations of all available major cations. In PNEC-pro, if Ca is not considered an input variable, the accuracy (adjusted *r*^*2*^) is less than 0.8 [5].

As a result, to apply a BLM-based risk assessment over a wider ecoregion, the major cations should be excluded from the minimum data requirements, and surrogate variables contributing to the good predictions for the BLM-based PNEC are required. In this study, electrical conductivity was considered a surrogate of the major cations and alkalinity. Electrical conductivity is typically included as a water quality variable in general regulatory water quality-monitoring databases. Electrical conductivity is one of the variables recommended for estimating the concentrations of missing BLM variables via its linear relationships with BLM variables [6, 7].

In the test datasets (four ecoregions), PNEC predictions were less reliable by the existing PNEC estimation tools than by the three different DNN models (Table 2). This was likely because the training datasets used for the development of each existing tool were not sufficiently representative of the different water chemistries, and the statistical and look-up table methods used for PNEC estimation provided limited predictive capacities for the nonlinear relationships between PNEC and BLM variables. Therefore, in this study, a training dataset representative of various freshwater chemistries was built for the DNN models. Its subsequent use resulted in a wide range of applications and good predictive capacity.

To design a representative training dataset, the frequencies of each BLM input variable and their relationships were investigated in the Korean freshwater monitoring database (S1 Fig). The domain ranges for water chemistry variables were determined from the abovementioned results (Fig 1). The pH conditions were generated as continuous values rather than multiple level conditions with intervals because pH was the only variable that had a non-linear relationship with PNEC. Another 9,792 combinations of Ca, Mg, Na, K, alkalinity, and DOC were generated assuming the same pH. Then 9,792 continuous pH variations were generated within the pH condition interval. These values were randomly arranged and added to the combined data of other variables.

The datasets used to develop the existing tools did not cover the full domain range of BLM input variables. For the mBAT training dataset, the Mg and Na concentrations and alkalinity were determined by Ca according to Peters et al. (2011) [9] and therefore consisted of a combination of only three variables: pH, DOC, and Ca. For the Bio-met training dataset, the Mg concentration was considered to be Ca-dependent, the Na concentration was considered to be dependent on four other factors, and alkalinity was determined to be dependent on pH as well as three other factors. The pH conditions of Bio-met were determined at 21 levels ranging from 6.0 to 8.5, while mBAT did not describe the pH conditions in detail. PNEC-pro, which was developed using monitoring data rather than simulation data, requires data from a wider ecoregion than just the Netherlands, the basis of its development.

To generate electrical conductivity data for the training dataset in this study, the use of MLR-based models to estimate electrical conductivity from BLM input variables has been proposed. To develop these models, the monitoring datasets from Korea, the United States, and Sweden were used because they included all BLM input variables and electrical conductivity. The final estimation model for electrical conductivity using Ca, Mg, and Na in Table 2 had a good predictive capacity, within a factor of two for 99.2% of the electrical conductivity data measured in the three ecoregions (*n* = 5,682) (S4 Fig). As a result, because the range of water chemistry data in the final training dataset with electrical conductivity covered the ranges of BLM input variables in the four test datasets (Korean, Swedish, United States, and Belgian freshwaters), it was considered to be sufficiently representative of the freshwater chemistry (Fig 3).

Better predictions of the copper PNECs were obtained from the three different types of DNN models trained and validated using the representative simulation training dataset than from the existing tools in the four test datasets (Korean, United States, Swedish, and Belgian freshwaters). The adjusted *r*^*2*^ values were higher than 0.95 in all but the Swedish freshwater dataset. Although the minimum adjusted *r*^*2*^ value in Swedish freshwater was 0.87, it was higher than the results obtained using the existing tools. The use of reduced input variables for the DNN(b) and DNN(c) models in Swedish freshwater, which had a lower pH and major cation concentration compared with the other regions, was probably why the adjusted *r*^*2*^ values (0.87 and 0.89, respectively) were lower than the value of 0.97 obtained with the DNN(a) model using all BLM variables (S2 Fig).

The mBAT and PNEC-pro predictions were less accurate than those of the DNN models, indicating that general statistical methods (multivariate polynomial regression and MLR) were not sufficient for predicting the nonlinear relationships between input variables and PNECs. A look-up table method, such as Bio-met, was expected to have a higher predictive capacity when used as the training dataset in this study, while the PNEC calculation performed in Excel required a considerable amount of time. The water chemistry conditions did not match the conditions in the training dataset, and its prediction accuracy was expected to be lower than that calculated by the DNN.

An important finding was the similar prediction accuracy in the test datasets of the three DNN models using different types of input variables to develop optimized PNEC estimation models depending on the available monitoring datasets. This means that even with reduced input variables, a good prediction capacity can be expected by a DNN model that includes the key input variables for a BLM. In particular, the DNN(c) model, which was selected as the most simplified surrogate tool, was shown to have a predictive capacity similar to that of the DNN(a) model, which provided the best prediction. Electrical conductivity played an important role as a variable acting as a surrogate for the major cations and alkalinity. Although there is further scope to reduce the uncertainty in the predicted PNECs by the DNN(c) model at a low pH and Ca concentration, such as in the Swedish freshwater, it is necessary to assess the environmental risk for copper using DNN(a) from all measured input variables. Consequently, according to the variables in the available monitoring databases, the most applicable model could be selected from among the three DNN models.

It is possible to reduce the uncertainty in the BLM-based PNECs estimated by the final surrogate tool in a specific region using a monitoring database containing the concentration of total organic carbon (TOC) rather than DOC. Both electrical conductivity and pH can be measured in field samples using commonly available water quality instruments and are included in most regulatory monitoring databases. The organic carbon concentration in freshwater is usually measured as TOC in monitoring databases unless the database is used for the purpose of bioavailability-based risk assessments. Among the test datasets in this study, the datasets from Korea, the United States, and Belgium included DOC concentrations for bioavailability-based risk assessments. The DOC concentration in the Swedish dataset was estimated by applying the 0.8 ratio, which is the simplest method of estimating DOC from TOC concentrations [11, 20]. However, the DOC concentration in Korean rivers is 64.3–79% of the TOC concentration, according to Kim et al. (2007) [21]. For surface waters in Poland and Germany, the DOC concentration range was 80–92% of the TOC concentration [22]. Thus, the observed DOC may be used to reduce the uncertainty of the BLM-based PNEC estimated using a surrogate tool.

## 5. Conclusion

This study developed three different types of DNN models, each requiring different input variables, which provide better predictions of the BLM-based PNECs for copper than existing PNEC tools in various ecoregions. The most applicable model among the three DNN models can be selected according to the available variables in monitoring databases. Furthermore, it is expected that the most simplified DNN model, using only general water quality variables (pH, DOC, and electrical conductivity), will enable the copper BLM-based risk assessment to be applied to monitoring datasets worldwide.

## Acknowledgments

This work was supported by Korea Environment Industry & Technology Institute (KEITI) through the Technology Development Project for Safety Management of Household Chemical Products funded by Korea Ministry of Environment (MOE) (2020002970009, 1485017560)

## Supporting information

S1 Fig. The relationships among biotic ligand model (BLM) input parameters and electrical conductivity within 764 samples from 93 sites in Korean freshwater.

S2 Fig. Comparison of the frequencies of biotic ligand model input variables in test datasets from United States, Korean, Swedish, and Belgian freshwaters.

S3 Fig. Comparison of the predicted no-effect concentrations (PNECs) from the multiple linear regression and biotic ligand model-based PNECs within the training dataset.

S4 Fig. Comparison of the measured electrical conductivity in the monitoring datasets (n = 5,682) from Korea, the United States, and Sweden with the electrical conductivity predicted by multiple linear regression.

S1 Table. Species- and element-specific parameters of chronic copper biotic ligand models.

S2 Table. The multiple linear regression formula for biotic ligand model variables for predicting electrical conductivity from Korean, Swedish, and United Sates monitoring databases.

## References

1. Di Toro DM, Allen HE, Bergman HL, Meyer JS, Paquin PR, Santore RC. Biotic ligand model of the acute toxicity of metals. 1. Technical basis. Environ Toxicol Chem. 2001; 20(10): 2383–96. https://doi.org/10.1002/etc.5620201034 PMID: 11596774

2. De Schamphelaere KA, Janssen CR. A biotic ligand model predicting acute copper toxicity for Daphnia magna: the effects of calcium, magnesium, sodium, potassium, and pH. Environ Sci Technol. 2002; 36(1):48–54. https://doi.org/10.1021/es000253sn PMID: 11817370

3. Bio-met. Bio-met Bioavailability Tool; UserGuide (Version5.0). 2019. https://bio-met.net/wp-content/uploads/2019/08/bio-met_Guidance-Document_v5.0_-2019-27-06.pdf

4. WFD UKTAG., Development and use of the copper bioavailability assessment tool (Draft). SC080021/8a-a; Water Framework Directive United Kingdom Technical Advisory Group: Scotland. 2012.

5. Verschoor AJ, Vink JP, Vijver MG. Simplification of biotic ligand models of Cu, Ni, and Zn by 1-, 2-, and 3-parameter transfer functions. Integr Environ Assess. and Manage. 2012; 8 (4):738–48. https://doi.org/10.1002/ieam.1298 PMID: 22556098

6. US EPA. Draft technical support document: recommended estimates for missing water quality parameters for application in EPA’s biotic ligand model, EPA 820-R-15-106; United States Environmental Protection Agency Office of Water 4304T: Washington DC. 2016.

7. McConaghie JB. Technical Support Document: An Evaluation to Derive Statewide Copper Criteria Using the Biotic Ligand Model, States of Oregon Department of Environmental Quality: Portland. 2016. https://doi.org/10.13140/RG.2.2.26803.32804

8. Thomas AJ, Petridis M, Walters SD, Gheytassi SM, Morgan RE. Two hidden layers are usually better than one. In: International conference on engineering applications of neural networks. Engineering Applications of Neural Networks. 2017; 279-290. https://doi.org/10.1007/978-3-319-65172-9_24

9. Peters A, Merrington G, De Schamphelaere KA, Delbeke K. Regulatory consideration of bioavailability for metals: Simplification of input parameters for the chronic copper biotic ligand model. Integr Environ Assess and Manage. 2011; 7(3):437–44. https://doi.org/10.1002/ieam.159 PMID: 21082669

10. De Schamphelaere KA, Heijerick DG, Janssen CR. Refinement and field validation of a biotic ligand model predicting acute copper toxicity to Daphnia magna. Comp Biochem Physiol Part C: Toxicol Pharmacol. 2002; 134:243–58. https://doi.org/10.1016/S1532-0456(02)00087-X PMID: 12356531

11. ECHA. Voluntary Risk Assessment of Copper, Copper II Sulphate Pentahydrate, Copper(I)Oxide, Copper(II)Oxide, Dicopper Chloride Trihydroxide. European Union Risk Assessment Report, European Copper Institute. 2008.

12. De Schamphelaere KA, Janssen CR. Development and field validation of a biotic ligand model predicting chronic copper toxicity to Daphnia magna. Environ Toxicol Chem. 2004; 23(6):1365–75. https://doi.org/10.1897/02-626 PMID: 15376521

13. Nys C, Regenmortel TV, Janssen CR, Oorts K, Smolders E, De Schamphelaere KA. A framework for ecological risk assessment of metal mixtures in aquatic systems. Environ Toxicol Chem. 2018; 37(3):623–42. https://doi.org/10.1002/etc.4039 PMID: 29135043

14. Kingma DP, Ba J. Adam: A method for stochastic optimization. CoRR. 2015; 1412.6980. https://arxiv.org/pdf/1412.6980.pdf

15. Nair V,Hinton GE, Rectified linear units improve restricted boltzmann machines. International Conference on Machine Learning: Proceedings of the 27th International Conference on International Conference on Machine Learning. Omnipress. 2010; 807-814. http://dblp.uni-trier.de/db/conf/icml/icml2010.html#NairH10

16. Schmidt-Hieber J. Nonparametric regression using deep neural networks with ReLU activation function. Ann Statist. 2020; 48 (4):1875–97. https://doi.org/10.1214/19-AOS1875

17. van Vlaardingen PL, Traas TP, Wintersen AM, Aldenberg T. ETX 2.0 A Program to Calculate Hazardous Concentrations and Fraction Affected, Based on Normally Distributed Toxicity Data, RIVM report 601501028; National Institute for Public Health and the Environment (RIVM): Bilthoven. 2004.

18. Natural Environment Research Council. Windermere Humic Aqueous Model; Use’s Guide (Version 7). Oxfordshire, UK. 2012.

19. US EPA. Aquatic Life Ambient Freshwater Quality Criteria-Copper, EPA-822-R-07-001; United States Environmental Protection Agency Office of Water 4304T: Washington DC. 2007.

20. Swedish Environmental Research Institute. Testing the biotic ligand model for Swedish surface water conditions - a pilot study to investigate the applicability of BLM in Sweden, IVL Report B1858; IVL Swedish Environmental Research Institute Ltd.: Stockholm. 2009.

21. Kim JK, Shin M, Jan C, Jung S, Kim B. Comparison of TOC and DOC distribution and the oxidation efficiency of BOD and COD in several reservoirs and rivers in the Han River system. J Korean Soc Water Environ. 2007; 23(1):72–80.

22. Sobczak P, Rosińska A. Concentration of total organic carbon and Its fractions in surface water in Poland and Germany. Proceedings. 2020; 51(1):35. https://doi.org/10.3390/proceedings2020051035

