## Supplement Figure 1 for "Development of a machine learning model to estimate biotic ligand model-based predicted no□effect concentrations for copper in freshwater"

**Supplementary material**

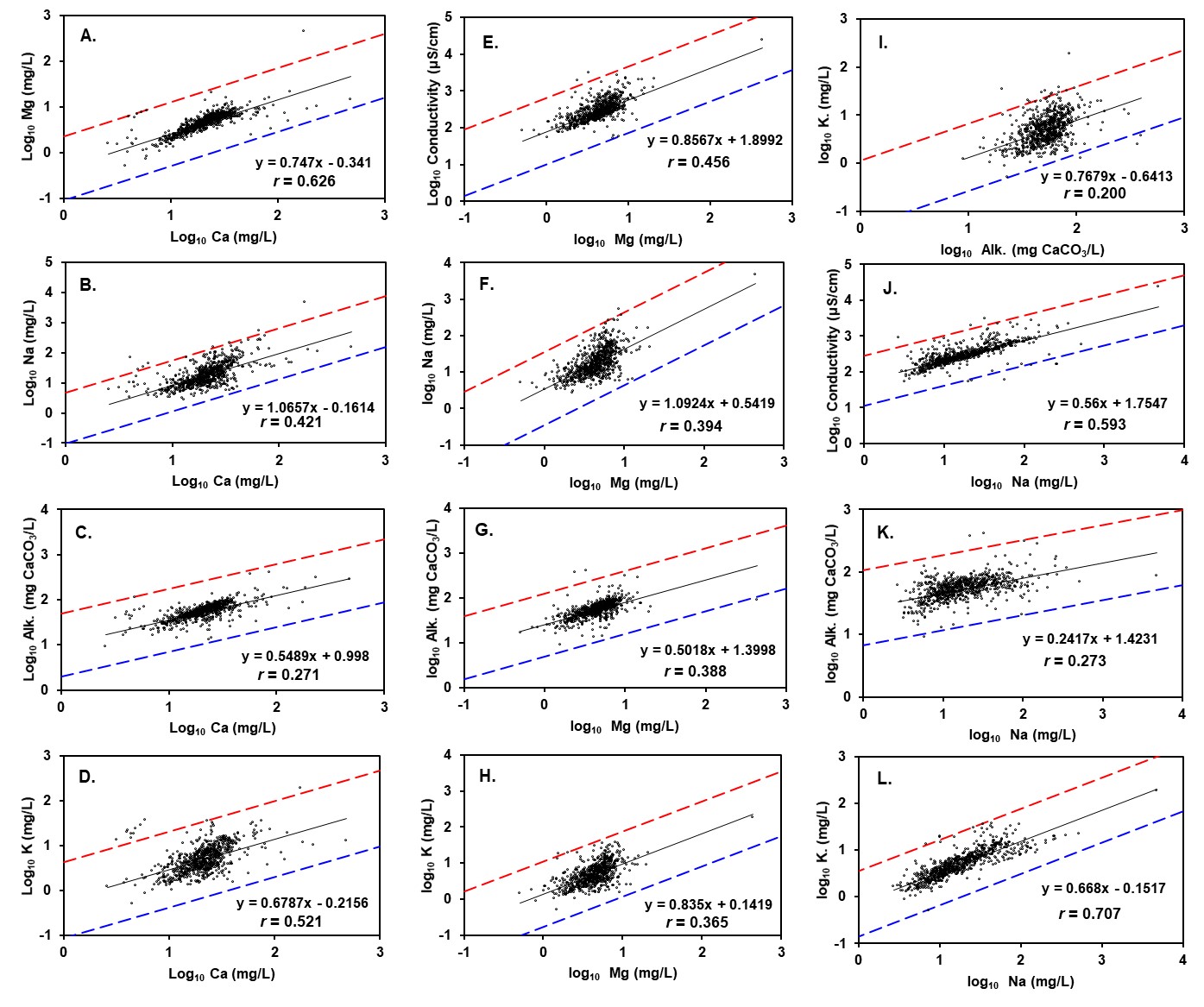

**S1 Fig.** The relationships among biotic ligand model (BLM) input parameters and electrical conductivity within 764 samples from 93 sites in Korean freshwater. The dashed lines indicate a factor of five from the linear regression line (solid line) for each BLM parameter (mg L^−1^ for cations; mg CaCO_3_ L^−1^ for alkalinity; µS cm^−1^ for electrical conductivity). *r* = correlation coefficient
