## Supplement Table 1 for "Development of a machine learning model to estimate biotic ligand model-based predicted no□effect concentrations for copper in freshwater"

**Supplementary material**

**S1 Table.** Species- and element-specific parameters of chronic copper biotic ligand models. pCu = -log of Cu^2+^ activity

| Species-specific parameter | *Fish* | *D. magna* | Algae / Plant |
| --- | --- | --- | --- |
| log K_CaBL_ | 3.47 | - |  |
| log K_MgBL_ | 3.58 | - |  |
| log K_NaBL_ | 3.19 | 2.91 | EC50_pCu_ |
| log K_HBL_ | 5.4 | 6.67 | = S_pH_ * pH + Q |
| log K_CuBL_ | 8.02 | 8.02 |  |
| log K_CuOHBL_ | 7.32 | 8.02 | S_pH_ = 1.354 ^1)^ / 1.136 ^2)^ |
| log K_CuCO3BL_ | 7.01 | 7.44 | Q = species-specific |
| R_CuOH_ = K_CuOHBL_/ K_CuBL_ | 0.2 | 1 | intercept |
| R_CuCO3_ = K_CuCO3BL_ / K_CuBL_ | 0.1 | 0.26 |  |
| Reference | De Schamphelaere *et al.* (2002) | De Schamphelaere and Janssen (2004) | ^1)^ De Schamphelaere *et al.* (2006); ^2)^ ECI, 2008 |
| Element-specific parameter |  |  |  |
| log K_CuCO3_ | 6.77 | | 6.75 |
| log K_Cu(CO3)2_^2-^ | 10.2 | | 9.92 |
| log K_CuHCO3_ | 12.13 | | 14.62 |
| Reference | Martell *et al*. (1997) | | Tipping E (1994) |
