## Supplement Figure 2 for "Development of a machine learning model to estimate biotic ligand model-based predicted no□effect concentrations for copper in freshwater"

**Supplementary material**

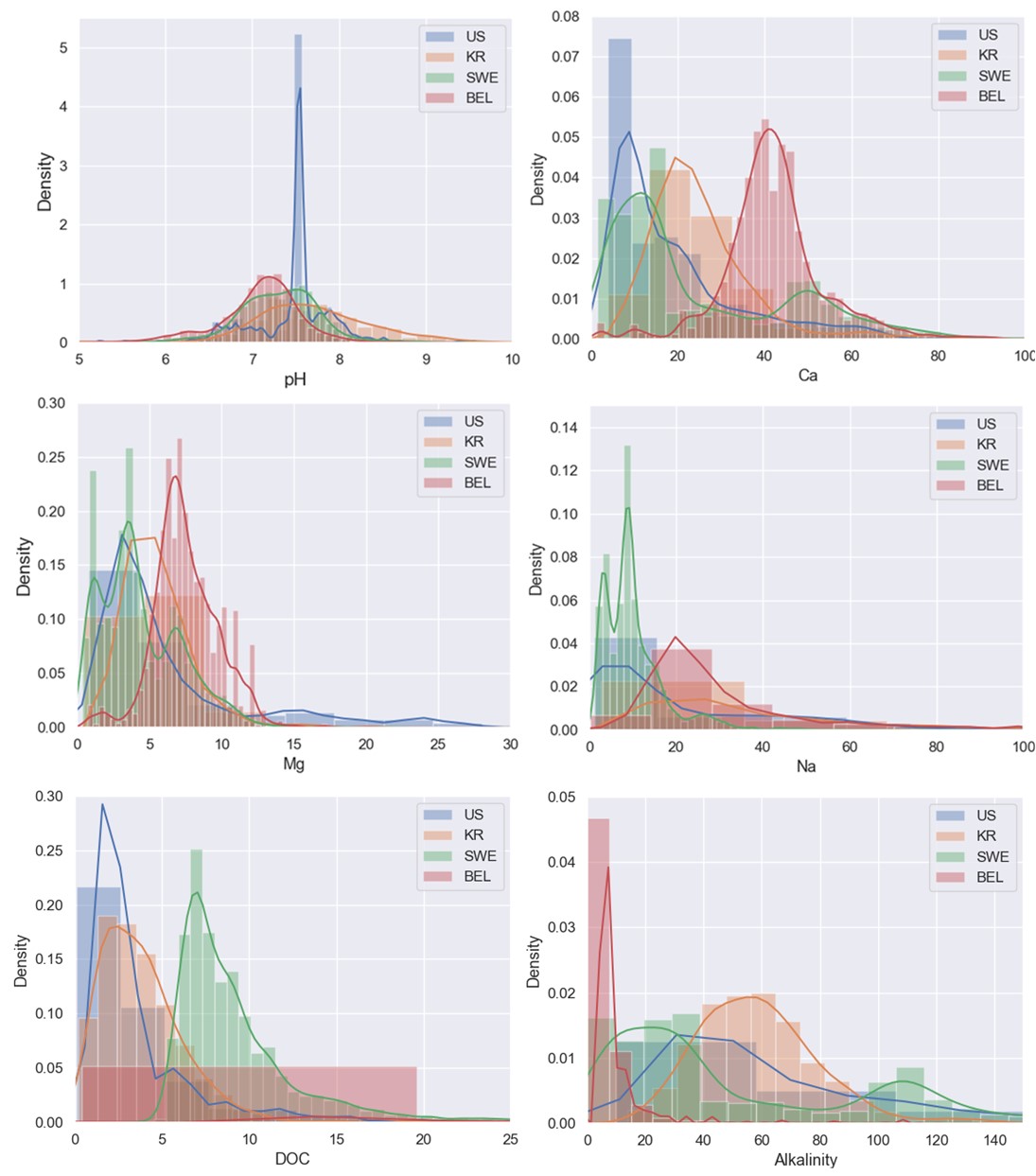

**S2 Fig.** Comparison of the frequencies of biotic ligand model input variables in test datasets from United States, Korean, Swedish, and Belgian freshwaters.
