## Supplement Table 2 for "Development of a machine learning model to estimate biotic ligand model-based predicted no□effect concentrations for copper in freshwater"

**Supplementary material**

**S2 Table.** The multiple linear regression formula for biotic ligand model variables for predicting electrical conductivity from Korean, Swedish, and United Sates monitoring databases with electrical conductivity

| N | Formula | Adj. *r^2^* | AIC | RSE | d.f. |
| --- | --- | --- | --- | --- | --- |
| 5 | 0.46 Ca+0.22 Mg+0.37 Na-0.02 Alkalinity-0.01 pH+1.29 | 0.9589 | −13914 | 0.07107 | 5676 |
| 4 | 0.46 Ca+0.22 Mg+0.37 Na-0.03 Alkalinity+1.23 | 0.9588 | −13900 | 0.07117 | 5677 |
| 3 | 0.43 Ca+0.22 Mg+0.36 Na+1.22 | 0.9587 | −13884 | 0.07127 | 5678 |

N = number of BLM variables; Adj. *r^2^* = adjusted *r^2^* value; AIC = Akaike information criterion; RSE = residual standard error; d.f. = degrees of freedom. Variables are log10 values (Ca, Mg, and Na in mg L^−1^, alkalinity in mg CaCO_3_ L^−1^) and pH.
