## Supplement Figure 3 for "Development of a machine learning model to estimate biotic ligand model-based predicted no□effect concentrations for copper in freshwater"

**Supplementary material**

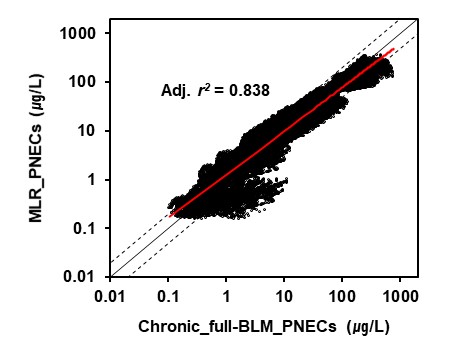

**S3 Fig.** Comparison of the predicted no-effect concentrations (PNECs) from the multiple linear regression and biotic ligand model-based PNECs within the training dataset. Adj. *r^2^* = adjusted *r^2^* value
