## Supplement Figure 4 for "Development of a machine learning model to estimate biotic ligand model-based predicted no□effect concentrations for copper in freshwater"

**Supplementary material**

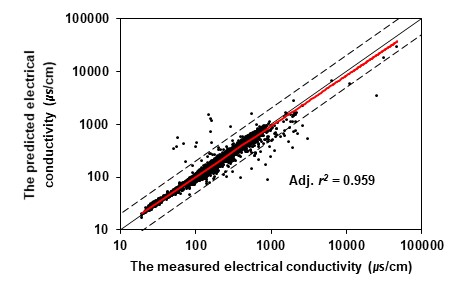

**S4 Fig.** Comparison of the measured electrical conductivity in the monitoring datasets (*n* = 5,682) from Korea, the United States, and Sweden with the electrical conductivity predicted by multiple linear regression. Adj. *r^2^* = adjusted *r^2^* value
